# Modulation of neuronal activity in cortical organoids with bioelectronic delivery of ions and neurotransmitters

**DOI:** 10.1101/2023.06.10.544416

**Authors:** Yunjeong Park, Sebastian Hernandez, Cristian O. Hernandez, Hunter E. Schweiger, Houpu Li, Kateryna Voitiuk, Harika Dechiraju, Nico Hawthorne, Elana M. Muzzy, John A. Selberg, Frederika N. Sullivan, Roberto Urcuyo, Sofie R. Salama, Elham Aslankoohi, Mircea Teodorescu, Mohammed A. Mostajo-Radji, Marco Rolandi

**Affiliations:** Department of Electrical and Computer Engineering, University of California Santa Cruz, Santa Cruz, CA 95064, USA; Genomics Institute, University of California Santa Cruz, Santa Cruz, CA 95060, USA; Live Cell Biotechnology Discovery Lab, University of California Santa Cruz, Santa Cruz, CA 95060; Centro de Electroquímica y Energía Química (CELEQ), Universidad de Costa Rica, San José, 11501 2060, Costa Rica; Department of Molecular, Cellular and Developmental Biology, University of California Santa Cruz, Santa Cruz, CA 95060

**Keywords:** bioelectronic ion pumps, ionic delivery, cortical organoids, stem cell models, organoid models

## Abstract

Precise modulation of brain activity is fundamental for the proper establishment and maturation of the cerebral cortex. To this end, cortical organoids are promising tools to study circuit formation and the underpinnings of neurodevelopmental disease. However, the ability to manipulate neuronal activity with high temporal resolution in brain organoids remains limited. To overcome this challenge, we introduce a bioelectronic approach to control cortical organoid activity with the selective delivery of ions and neurotransmitters. Using this approach, we sequentially increased and decreased neuronal activity in brain organoids with the bioelectronic delivery of potassium ions (K^+^) and γ-aminobutyric acid (GABA), respectively, while simultaneously monitoring network activity. This works highlights bioelectronic ion pumps as tools for high-resolution temporal control of brain organoid activity toward precise pharmacological studies that can improve our understanding of neuronal function.

## INTRODUCTION

Pluripotent stem cells-derived cortical organoids are valuable tools to study brain development, evolution, and disease ^1,2^. Several efforts have tried to understand the emergence, development, and maturation of circuit formation in these microphysiological systems ^3-6^. Moreover, circuit activity manipulation by either grafting of defined cell types ^7,8^ or treatment with small molecules ^9,10^ has helped dissect the cellular and molecular mechanisms fundamental characteristics of neurodevelopmental disorders. During development, neuronal activity plays a key role in cortical circuit establishment and maturation ^11-13^. For example, neuronal activity modulation controls several processes, including fate acquisition and neuronal migration ^14-16^. Similarly, other molecules, including neurotransmitters, regulate the proliferation of radial glia cells and control the migration of interneurons to the cerebral cortex ^17-21^. Despite the important roles of ions and neurotransmitters in cortical development, current organoid models lack the temporal resolution to study subtle changes in the concentrations of these ions and molecules in discrete processes. Scientists either use cell lines mutated for the receptors of interest ^8^ or perform bath applications of the desired ions and molecules ^22,23^. To address these challenges, bioelectronic electrophoretic pumps deliver ions and charged molecules with high spatiotemporal control ^24-26^. Bioelectronic ion pumps offer distinct advantages, including hydrogel-based selectivity toward specific ions, reduced invasiveness, and the ability to perform continuous and automated ion delivery ^27-29^. Moreover, the development of multi-ion pumps can allow the simultaneous or sequential control of several processes ^30,31^. Previous work has demonstrated the efficacy of these pumps in *in vivo* models of inflammation ^32^, wound healing ^33^ and epilepsy ^34^, as well as the delivery of potassium ions to *in vitro* 2D models to actuate and control biological processes ^24^. However, their applications to organoids and other *in vitro* 3D systems remain largely unexplored. Here, we have demonstrates the modulation of organoid neuronal activity with the spatiotemporal delivery of K^+^ ions and small molecules γ-aminobutyric acid (GABA) to brain organoids with bioelectronic ion pumps. (Figure 1).

**Figure 1.**
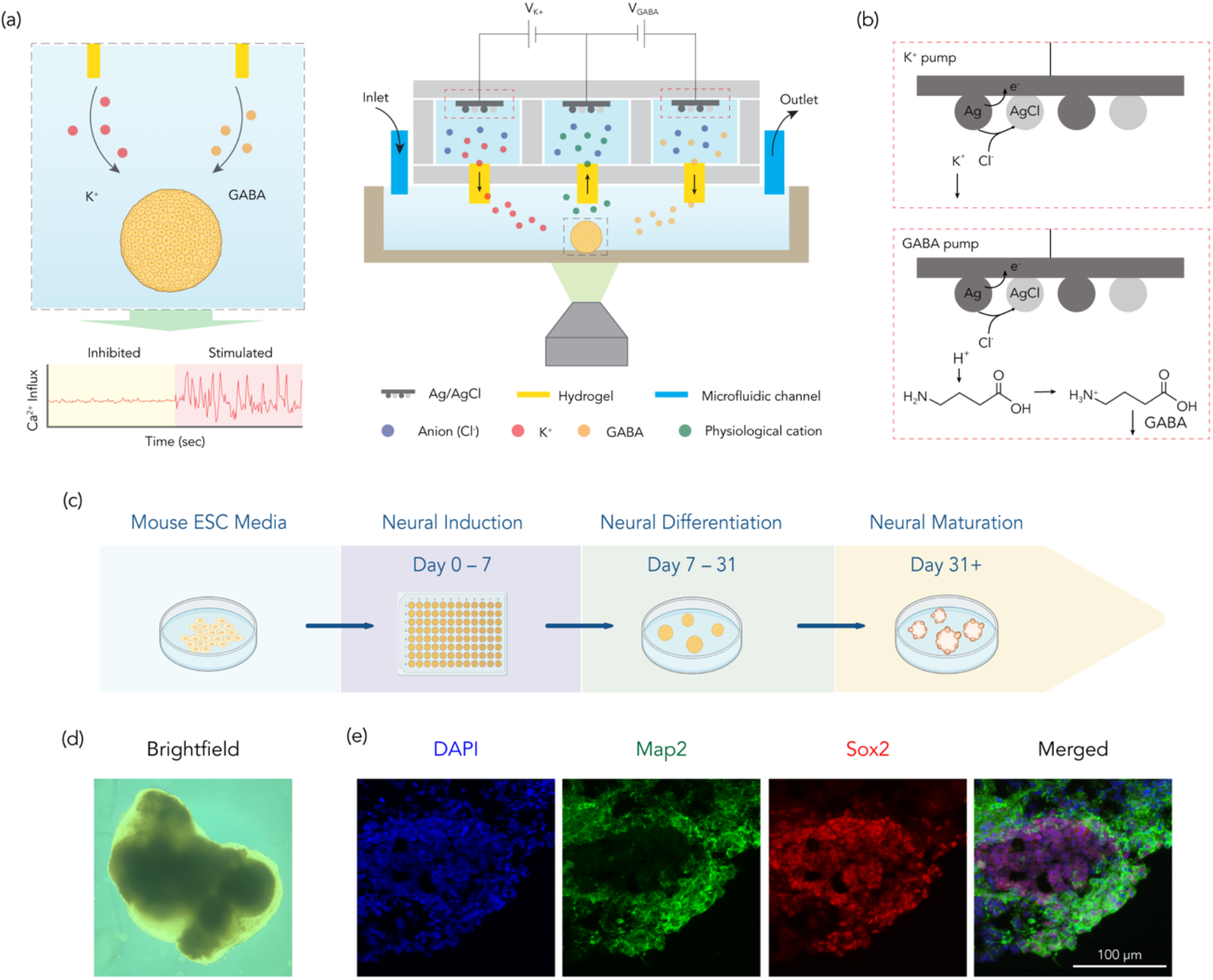
Experimental Design. (a) (left) Schematic representation of the bioelectronic ion pump for targeted delivery of K^+^ and GABA. This is combined with a fluorescence microscope that enables real-time monitoring of the organoids’ activity via calcium imaging. (right) A detailed schematic representation of the ion pump mechanism, illustrating the direction of ion movement during operation. (b) Electrochemical reaction at the working electrode with (top) K^+^ and (bottom) GABA molecules. (c) Schematic representing the protocol for cortical organoid generation and culture (d) Bright-field image of a cortical organoid. (e) Immunostaining of a cortical organoid showing the presence of neurons (Map2; green) and neuronal progenitors (Sox2; red). Nuclear counterstain is performed with DAPI.

## RESULTS

### Bioelectronic delivery of K^+^ and GABA

As previously described, bioelectronic ion pumps have demonstrated the capacity to deliver ions and charged molecules to various *in vivo* and *in vitro* 2D models ^24,30^. However, their application in organoids and other 3D *in vitro* systems has remained largely uncharted. To extend this technology, we adapted a specialized bioelectronic ion pump intended for delivery within 3D organoid models (Figure 1a, left). This ion pump delivers GABA and K^+^ ions, either individually or concurrently, to the surface of mouse cortical organoids. As both GABA, a neurotransmitter, and K^+^, a modulator of neural excitability, play vital roles in neural activity ^35,36^, their delivery offers the opportunity to observe the combined effects of these charged molecules on intracellular calcium dynamics, a reliable marker of neural activity (Figure 1a, left).

To facilitate the integration of the ion pump with *in vitro* cultures, we designed a 3D-printed adapter (Figure S1). This adaptor stabilizes the ion pump, allowing a snug fit into a standard six-well cell culture plate (Figure S1). This adapter design enables media exchange without displacing the cortical organoid, ensuring stable experimental conditions. The bioelectronic pump contains three chambers, two of which hold 1 M KCl and 100 mM GABA solutions, along with a spare chamber that contains the reference electrode (Figure 1a, right). With a potential difference between the working electrode (WE) and reference electrode (RE) (V_K^+^_ for K^+^ delivery and V_GABA_ for GABA delivery), the pump delivers K^+^ and GABA from the reservoirs to the target (Figure 1a, right).

In the K^+^ delivery system, a V_K^+^_ prompts the WE to attract Cl^-^ ions. These ions undergo oxidation on reaching the Ag surfaces, resulting in the formation of AgCl (Figure 1b, top). The electric field then drives the resulting K^+^ ions from the WE-containing reservoir into the target through the capillary (Figure 1b, top). The V_K^+^_ could potentially prompt physiological cations, such as Na^+^ or Ca^2+^, to move back into the spare chamber containing the RE ^24,30^. In contrast, the ion pump effectively transports K^+^ at high concentrations, which created a more pronounced effect on the organoid ^24,30^.

For GABA delivery, adjusting the pH of the solution to 4 with HCl enables protonation of the GABA molecules, leading to the formation of positively charged γ-aminobutyric acid cations ^34^. These cations have the potential to move through the anionic hydrogel-filled capillary under the influence of an electric field ^37^. The V_GABA_ between the WE and RE attracts Cl^-^ ions to the Ag surfaces of the WE, producing AgCl in a process similar to the K^+^ delivery process (Figure 1b, bottom). Subsequently, the protonated GABA molecules in the reservoir then acquired a positive charge and responded to the electric field generated by the WE. These positively charged GABA molecules traveled through the capillary, which selectively transported cations between the reservoir and the target (Figure 1b, bottom).

To test the applicability of bioelectronic ion pumps to modulate organoids, we generated cortical organoids using mouse embryonic stem cells (ESCs). We modified an established protocol ^38,39^ to optimize the maturation of neurons (Figures 1c–d). We validated the presence of neuronal progenitors and neurons in these organoids by performing immunohistochemical staining for Sox2 and Map2, respectively (Figure 1e). Together, these bioengineered approaches allow us to test the action of ionic pumps in 3D organoid systems.

### K^+^ delivery using bioelectronic ionic pumps leads to rapid excitation of neuronal cells in cortical organoids

We set up the well plate with the 3D holder over a fluorescence microscope to demonstrate and monitor K^+^ delivery from the ion pump to the cortical organoids in real time. Before deliver the K^+^, we characterized the ion pump delivery of K^+^ ions, we monitored the fluorescence intensity of a K^+^ indicator, Ion Potassium Green-2, a dye that linearly changes its fluorescence intensity in response to variations in K^+^ concentration. We maintained alternating V_K^+^_ pulses of +1 V and -1 V for 1 minute over five cycles. The V_K^+^_ = +1 V delivered K^+^ ions from the reservoir to the target, while the V_K^+^_ = -1 V moved cations in the opposite direction. Following the calibration of fluorescent response against known K^+^ ion concentrations (Figure S2a), we found that the ion pump could deliver 3 mM of K^+^ ions to the target during a 1-minute actuation period (Figure 2b). To evaluate the effect of the bioelectronic ion pump on cortical organoid neuronal activity, we conducted calcium imaging on a region of interest (ROI) encompassing three stages. This ROI, demarcated by a white dotted line, is shown in Figure 2c. Each stage had corresponding images taken at key intervals to further illustrate the process (Figure 2c, left). Notably, black arrows on the right plot highlight the specific times of image capture, allowing for a temporal correlation with the observed effects (Figure 2c). In the first stage of the calcium imaging, we established baseline conditions by recording the inherent spontaneous neuronal activity of the cortical organoids before the introduction of any ions via the bioelectronic ion pump. Changes in fluorescence, representing calcium transients in the ROIs, signified the spontaneous activity (Figure 2c; images i and ii). This stage played a crucial role in providing a comparison point between the activation and deactivation stages of the bioelectronic ion pump. While all cortical organoids exhibited blinking spots indicative of calcium mobilization into cells; each organoid showed different calcium responses, as illustrated in Figure 2c. For example, the first organoid exhibited relative quietness with some noise, while the third organoid presented irregular and comparatively faster transitions in calcium dynamics.

**Figure 2.**
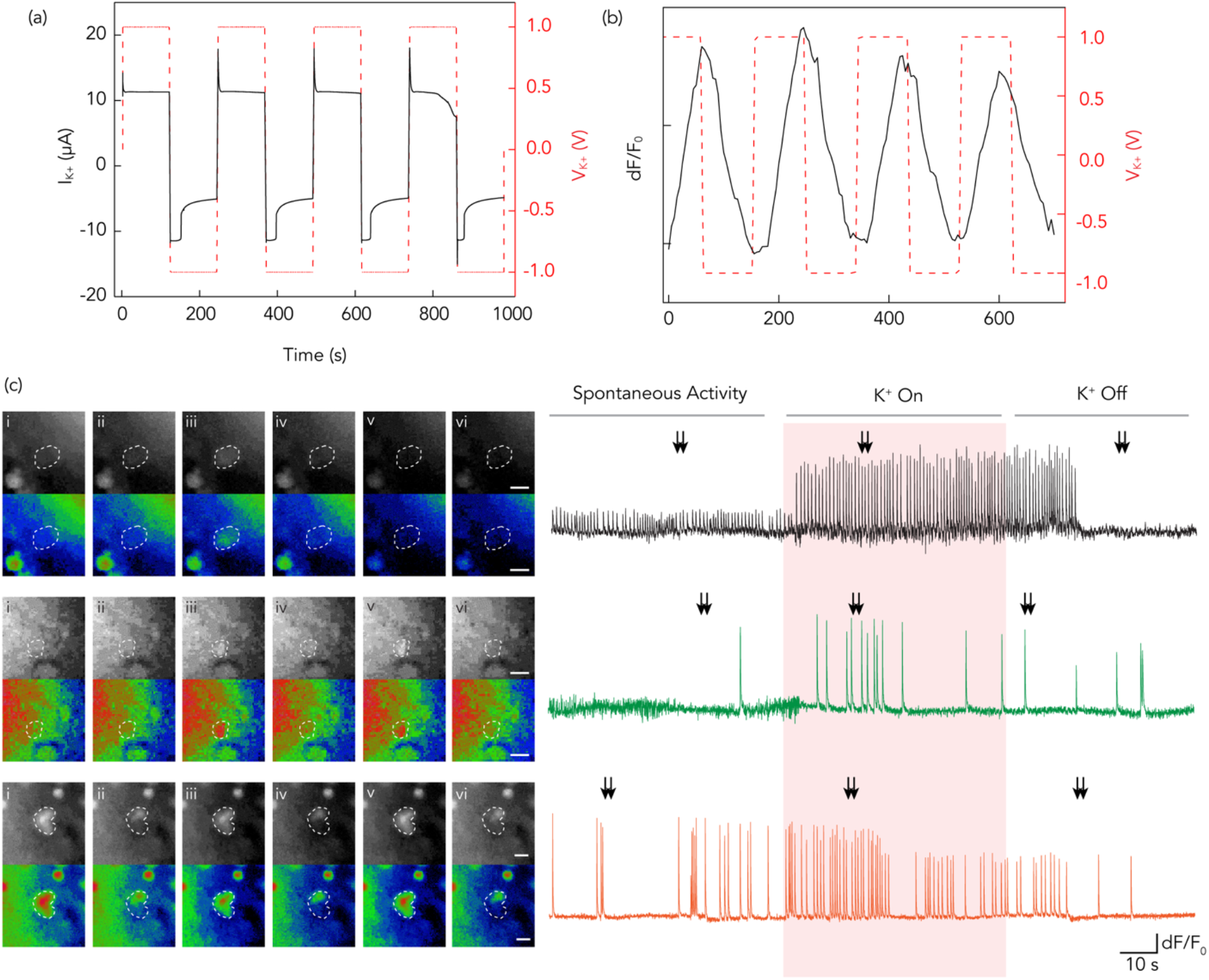
Bioelectronic delivery of K^+^ ions to cortical organoids. (a) Electrical characterization of the bioelectronic ion pump, depicting the current response (I_K^+^_) of the bioelectronic K^+^ ion pump according to the V_K^+^_. (b) Fluorescence response of bioelectronic K^+^ ion pump in accordance with V_K^+^_. (c) Calcium imaging of organoids: (left) Representative images of calcium imaging at distinct stages of neuronal activity correlating with the time points indicated by the black arrows on the right plot. (right) A plot of fluorescence intensity versus time, showcasing the dynamic changes in neuronal activity during the K^+^ ion delivery process. (Scale bar: 10 μm)

Proceeding the second stage, the activating the bioelectronic ion pump (K^+^ On) led to a significant increase in calcium mobilization (Figure 2c; images iii and iv). The augmentation of K^+^ ions crucially alter the ionic environment, triggering the excitatory effect on the cortical organoid, thus explaining this observable increase ^35,40^. Within ten seconds of ion pump activation, the first organoid exhibited calcium transients, maintaining a nearly constant rate of signal frequency throughout the period of measurement. Similarly, an accelerated signal frequency ensued post ion pump activation for the second organoid, although the calcium transients revealed instability, exhibiting an erratic surge. The third organoid responded with a rapid increase in calcium signaling after K^+^ delivery.

Transitioning to the third stage, following the deactivation of the ion pump (K^+^ Off) and subsequent wash with fresh media, we observe a decline in the activity levels of all organoids. A slower signal frequency suggested a reversible action induced by ion delivery from the system (Figure 2c; images v and vi). The first organoid persisted in a stimulated state after washing, eventually reverting to a quieter state. The second organoid exhibited a slower response compared to the stimulated state, with irregular fluorescence intensity and frequency of calcium transients. Similarly, the third organoid, after a certain duration, underwent a gradual transition to a quieter state, similar to the first organoid’s behavior. These observations collectively established that K^+^ ions could effectively stimulate cortical organoids, exhibiting a reversible effect.

### Bioelectronic pumps can effectively inhibit neuronal activity in brain organoids by delivering GABA

We set up the inhibitory neurotransmitter of the GABA delivery ion pump. To accurately quantify the amount of GABA delivered, we employed liquid chromatography (LC) as a sensitive and reliable analytical technique. Similar to the K^+^ delivery experiment, the experimental protocol involved alternating V_GABA_ pulses at +1 V and -1 V, each maintained for one minute. The V_GABA_ = 1 V pushes GABA from the reservoir to the target, while the V_GABA_ = -1 V moves ions in the opposite direction. We recorded the current (I_GABA_) generated upon V_GABA_ and presented the data in Figure 3a.

**Figure 3.**
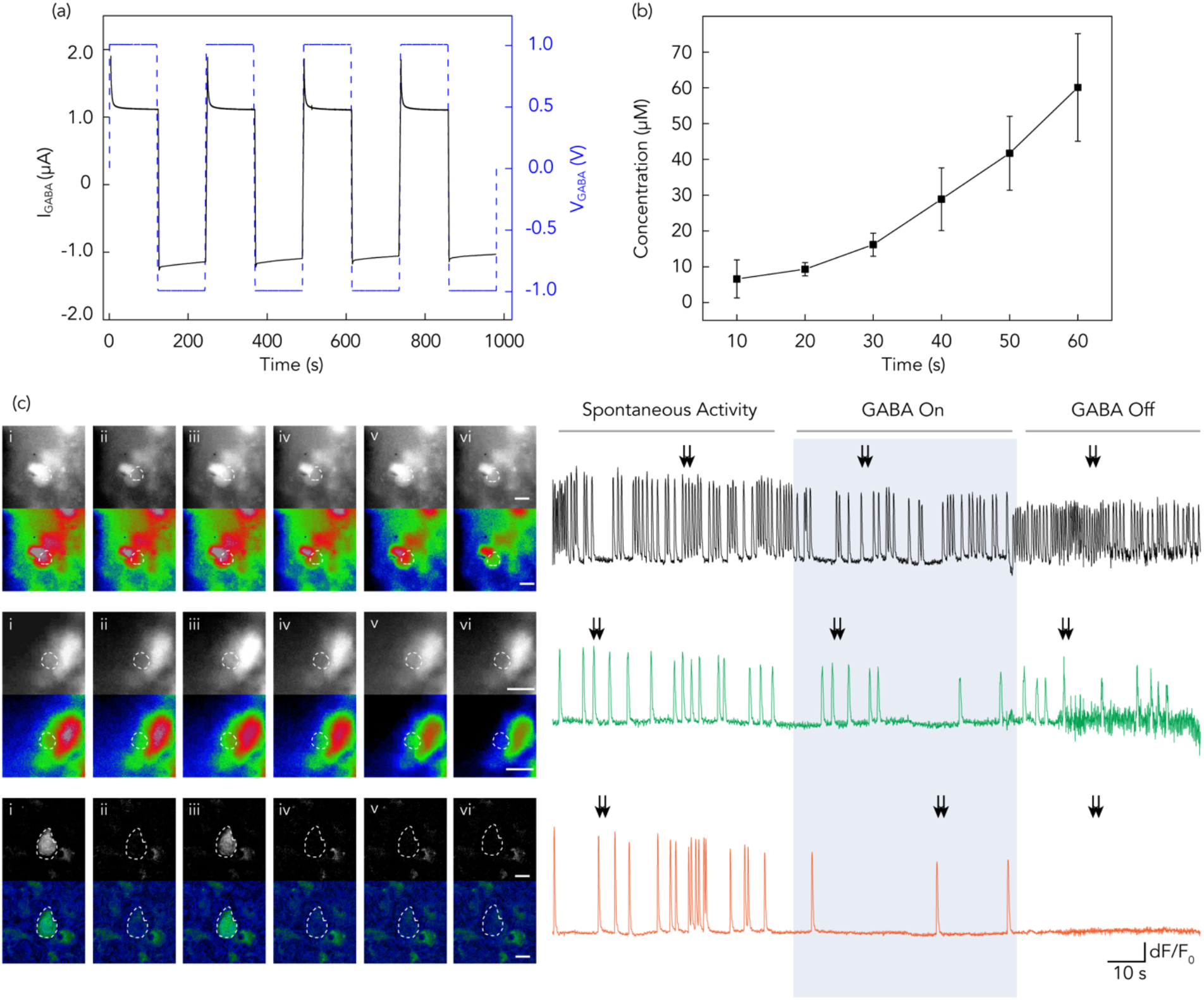
Bioelectronic delivery of GABA to cortical organoids. (a) Electrical characterization of the bioelectronic GABA ion pump, depicting the current response (I_GABA_) of the bioelectronic GABA ion pump according to the V_GABA_. (b) Liquid chromatography (LC) analysis of GABA concentration, demonstrating a detectable increase in concentration over a 1-minute delivery period in response to the V_GABA_. (c) Calcium imaging of organoids: (left) Representative images of calcium imaging at distinct stages of neuronal activity corresponding to the time points indicated by the black arrow on the right plot. (right) A plot of fluorescence intensity versus time, illustrating the dynamic changes in neuronal activity during the GABA delivery process. (Scale bar: 10 μm)

After the delivery process, we collected samples from the target and performed LC measurements to measure the GABA concentration. The result presented in Figure 3b illustrates a clear, time-dependent increase in the GABA concentration within the target, demonstrating the effective delivery of GABA via our ion pump. Calibration data, cross-verified against known GABA concentration, confirmed the delivery performance of the ion pump and indicated the delivery of approximately 70 μM of GABA within 1 minute actuation period (Figure 3b and Figure S2b).

Using the previous procedure with K^+^ ions as a reference method, we introduced GABA as an effector. We then conducted calcium imaging on an ROI within the cortical organoids. As displayed in Figure 3c, all cortical organoids within the ROI revealed blinking spots, signifying varied calcium dynamics. The first organoid exhibited rapid calcium transients, whereas the second and third organoids showed comparable slower signaling patterns (Figure3c; image i and ii). Upon activating the pump (GABA On), we noted a considerable decline in neural activity, indicated by reduced blinking frequency (Figure 3c; image iii and iv). Each organoid measured within the ROI exhibited a decrease in calcium mobilization, thus entering a notably quieter state. Following the deactivation of the ion pump and washing the organoid with fresh media (GABA Off), we noted a resurgence in neural activity within the ROI, exhibited by the resumption of blinking or the increase in overall activity (Figure 3c). Despite this, blinking frequencies recovered at different ratios. For instance, the third organoid did not return to its original state after washing with fresh media due to varying organoid resilience and responsiveness to ion modulation, or pre-existing conditions affecting its recovery.

### Sequential GABA and K^+^ delivery can rapidly modulate organoid activity

To investigate the combined effects of GABA and K^+^ ions on neural activity in cortical organoids, we structured an experiment encompassing a series of phases (Figure 4). Initially, within a defined ROI, we observed and recorded the intrinsic neural activity of the organoids without any stimulations to establish a spontaneous activity baseline as described before (Figure 4; images i and ii). Next, we activated the bioelectronic ion pump to deliver GABA (GABA On) to the organoids, while real-time monitoring captured the evolving neural activity within the ROI. The resulting effect exhibited an expected inhibitory response in the cortical organoids (Figure 4; images iii – vi) After deactivating the GABA device (GABA Off), the organoid underwent a fresh media wash, which enabled the removal of delivered GABA and facilitated a transition of the organoid back towards their original condition (Figure 4; images vii – x). We then activated the ion pump once again (K^+^ On), this time for K^+^ ion delivery to the organoids, while consistently tracking the resultant neural activity within the ROI. In the final phase, we deactivated the K^+^ ion pump (K^+^ Off) and performed another round of fresh media wash on the organoids to remove the delivered K^+^, allowing them to return to a state closer to their original condition.

**Figure 4.**
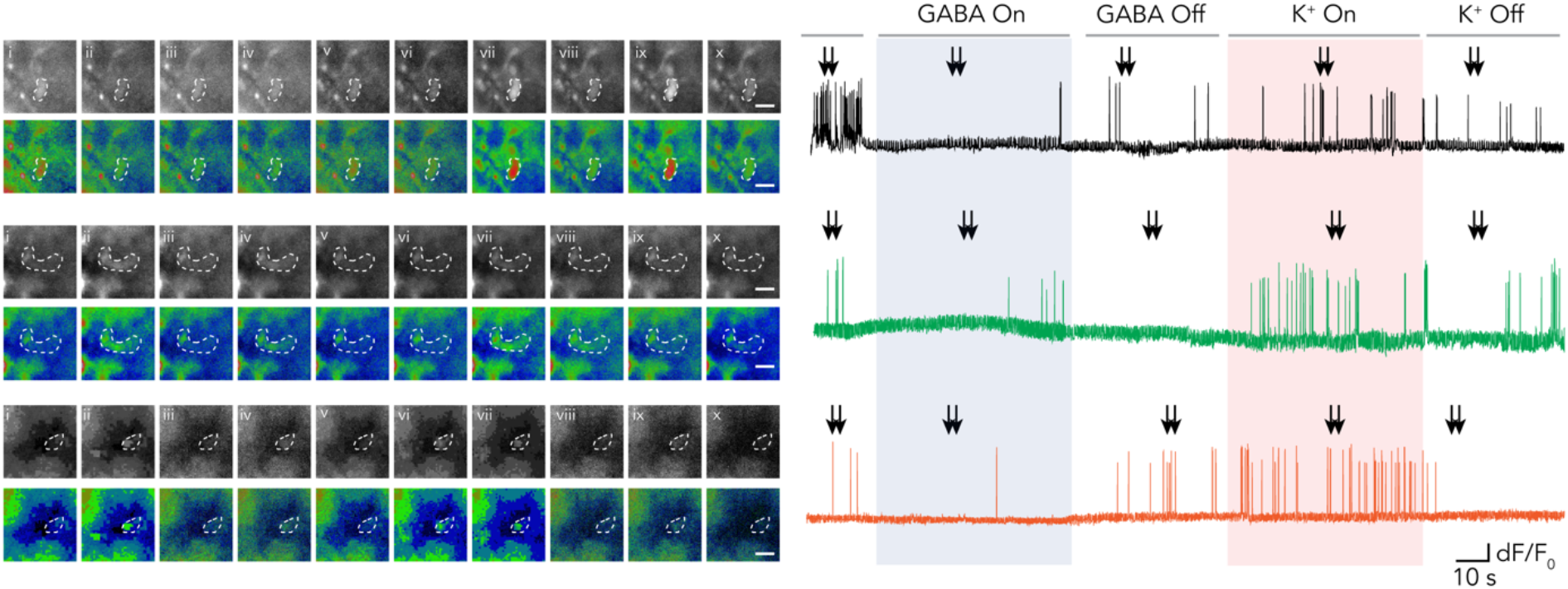
Sequential modulation of neural activity in cortical organoids using GABA and K^+^ ions. (left) Representative calcium imaging images capture distinct stages of neuronal activity in organoids following the sequential delivery of GABA and K^+^ using the bioelectronic ion pumps corresponding to the time points indicated by the black arrow on the right plot. (right) The changes in fluorescence intensity over time, show the dynamic changes in neuronal activity. An initial decrease in blinking frequency highlights the inhibitory effect of GABA delivery, succeeded by an increase in blinking frequency signifying the excitatory effect of K^+^ delivery. (Scale bar: 10 μm)

The series of experiments on the cortical organoids, offered valuable insights into the individual and combined impacts of GABA and K^+^ ion delivery, facilitated by the bioelectronic ion pump. The studies found that the GABA delivery initiated a significant decrease in neuronal activity, while the following delivery of K^+^ ions further modulated neural activity, leading to an elevated activity state. The effects of both ion deliveries proved reversible, as indicated by the gradual return of the organoids within the ROI to an original state or quieter state after washing away the delivered ions. This study highlights the potential of bioelectronic devices to enable controlled and reversible modulation of neural activity.

### Conclusions

The ability to rapidly manipulate organoid activity can have direct applications in both fundamental and translational biology studies ^24,26,41^. In this work, we have demonstrated the application of a bioelectronic ion pump to increase and inhibit neuronal activity with the targeted delivery of K^+^ and GABA to cortical organoids. Using calcium transients as the readout for baseline neuronal activity, we delivered K^+^ ions to increase activity and GABA to inhibit neuronal activity. We observed that the organoids returned to their initial state upon washing with fresh media, indicating the potential for reversible experiments. This work marks a significant advancement over current approaches, which often introduce pharmacological agents through bath applications requiring lengthy incubation periods for agent diffusion ^5^. Considering the rapid return to the baseline state, these pumps could facilitate chronic electrophysiology experiments, currently limited to recording the spontaneous activity of the organoids ^4,42^. Future applications could include combining these pumps with high-density multielectrode arrays ^4^ which will open new avenues to study circuit maturation and plasticity.

## MATERIALS AND METHODS

### ESC culture

All experiments were performed in the ES-E14TG2a mouse ESC line (ATCC CRL-1821). This line is a derived from a male of the 129/Ola mouse strain. Mycoplasma testing confirmed lack of contamination.

ESCs were maintained on Recombinant Human Protein Vitronectin (Thermo Fischer # A14700) coated plates using mESC maintenance media containing Glasgow Minimum Essential Medium (Thermo Fisher Scientific # 11710035), Embryonic Stem Cell-Qualified Fetal Bovine Serum (Thermo Fisher Scientific # 10439001), 0.1 mM MEM Non-Essential Amino Acids (Thermo Fisher Scientific # 11140050), 1 mM Sodium Pyruvate (Millipore Sigma # S8636), 2 mM Glutamax supplement (Thermo Fisher Scientific # 35050061), 0.1 mM 2-Mercaptoethanol (Millipore Sigma # M3148), and 0.05 mg/ml Primocin (Invitrogen # ant-pm-05). mESC maintenance media was supplemented with 1,000 units/mL of Recombinant Mouse Leukemia Inhibitory Factor (Millipore Sigma # ESG1107). Media was changed daily.

Vitronectin coating was incubated for 15 min at a concentration of 0.5 μg/mL dissolved in 1X Phosphate-buffered saline (PBS) pH 7.4 (Thermo Fisher Scientific # 70011044). Dissociation and cell passages were done using ReLeSR passaging reagent (Stem Cell Technologies # 05872) according to manufacturer’s instructions. Cell freezing was done in mFreSR cryopreservation medium (Stem Cell Technologies # 05855) according to manufacturer’s instructions.

### Cortical organoids generation

To generate cortical organoids we clump-dissociated ESCs using ReLeSR and re-aggregated in lipidure-coated 96-well V-bottom plates at a density of 10,000 cells per aggregate, in 200 μL of mESC maintenance media supplemented with Rho Kinase Inhibitor (Y-27632, 10 μM, Tocris # 1254) (Day -1). After one day (Day 0), we replaced the medium with cortical differentiation medium containing Glasgow Minimum Essential Medium (Thermo Fisher Scientific # 11710035), 10% Knockout Serum Replacement (Thermo Fisher Scientific # 10828028), 0.1 mM MEM Non-Essential Amino Acids (Thermo Fisher Scientific # 11140050), 1 mM Sodium Pyruvate (Millipore Sigma # S8636), 2 mM Glutamax supplement (Thermo Fisher Scientific # 35050061) 0.1 mM 2-Mercaptoethanol (Millipore Sigma # M3148) and 0.05 mg/ml Primocin (Invitrogen # ant-pm-05).

Cortical differentiation medium was supplemented with Rho Kinase Inhibitor (Y-27632, 20 μM # 1254), WNT inhibitor (IWR1-ε, 3 μM, Cayman Chemical # 13659) and TGF-Beta inhibitor (SB431542, Tocris # 1614, 5 μM, days 0-7). Media was changed on days 3 and 6 and then every 2-3 days until day 7. On day 7 organoids were transferred to ultra-low adhesion plates (Millipore Sigma # CLS3471) and put on an orbital shaker in neuronal differentiation medium at 75 revolutions per minute. Neuronal differentiation medium contained Dulbecco’s Modified Eagle Medium: Nutrient Mixture F-12 with GlutaMAX supplement (Thermo Fisher Scientific # 10565018), 1X N-2 Supplement (Thermo Fisher Scientific # 17502048), 1X Chemically Defined Lipid Concentrate (Thermo Fisher Scientific # 11905031) and 0.05 mg/ml Primocin (Invitrogen # ant-pm-05). Organoids were grown under 40% O_2_ and 5% CO_2_ conditions. Medium was changed every 2-3 days.

On day 14 onward, we added 5 μg/mL Heparin sodium salt from porcine intestinal mucosa (Millipore Sigma # H3149) and 0.5% v/v Matrigel Growth Factor Reduced (GFR) Basement Membrane Matrix, LDEV-free (Matrigel GFR, Corning # 354230) to the neuronal differentiation medium.

On day 21 onward, we transferred the organoids to neuronal maturation media containing BrainPhys Neuronal Medium (Stem Cell Technologies # 05790), 1X N-2 Supplement, 1X Chemically Defined Lipid Concentrate (Thermo Fisher Scientific # 11905031), 1X B-27 Supplement (Thermo Fisher Scientific# 17504044), 0.05 mg/ml Primocin (Invitrogen # ant-pm-05). and 1% v/v Matrigel Growth Factor Reduced (GFR) Basement Membrane Matrix, LDEV-free.

### Fabrication of ion pump

Fabricating an ion pump involved a combination of a PDMS reservoir, hydrogel capillaries, and a PCB control board. Formlabs 3D printers were used to print the bottom and top layers of the reservoir. Ag and AgCl wires were inserted into the reservoirs to create the electrodes, and the two parts of PDMS were bonded after being treated with 50 W oxygen plasma for 10 s. A 1.5μm thick water-insulating layer of parylene-C was deposited using the Coating Systems Lab coater two systems in the presence of an A174 adhesion promoter. Four-7mm long hydrogel-filled capillaries through the bonded PDMS and the reservoirs were filled with 1M KCl solution using a syringe. The PCB board was soldered onto the PDMS device and connected using silver pastes and ally dowel pins.

The hydrogel in the capillaries consisted of 2-Acrylamido-2-methylpropane sulfonic acid (AMPSA) and Poly (ethylene glycol) diacrylate (PEGDA). The glass capillaries were treated with Silane A-174 (3-(Trimethoxysilyl) propyl methacrylate) and crosslinked under UV light for 5 min. The control board was a 16-channel potentiostat interfaced to a Raspberry Pi single-board computer. The Raspberry Pi was put into soft Access Point (AP) mode for a laptop connected to it via Wi-Fi and provided necessary actions based on the current readings. The ring-shaped PCB board featured four plated through holes, which served as the insertion points for dowel pins. These pins, in turn, were connected to the control board through a JST-SH 4-pin connector, establishing a secure and reliable electrical connection between the four reservoirs and the control board.

### Device characterization

To monitor variation in K^+^ ion and GABA concentration, we employed micro-based real-time imaging and LC measurements. For K^+^ ions, we utilized IPG-2 dye, an intracellular K^+^ indicator optimized to a concentration of 3 μM in 0.1M Tris buffer. This dye, characterized by excitation and emission wavelengths of 525 nm and 545 nm respectively, exhibiting a linear relationship between fluorescence intensity and K^+^ concentration, thereby enabling the detection of subtle shifts in K^+^ levels. The data collected were analyzed using ImageJ 1.53p version. For GABA quantification, LC measurement was carried out using LC-MS (LTQ-Orbitrap, Thermo, USA)

### Calcium imaging

The day before imaging, the organoids were switched to maturation media in which BrainPhys Neuronal Media was replaced with BrainPhys Imaging Optimized Medium (Stem Cell Technologies # 5796). Calcium imaging was done using 4μM Fluo-8 AM (Abcam # ab142773). The organoids were incubated at 37 °C with Fluo-8 AM for 30 minutes before imaging. All imaging was performed using an EVOS M7000 Imaging System with images taken every (40 frames per second) and processed using ImageJ 1.53p version. Only organoids with baseline calcium transients were used for imaging experiments.

### Ion delivery using the bioelectronic ion pump

The bioelectronic ion pump was integrated with organoids placed in a six-well plate for the ion delivery experiments. In our in vitro experiments, we used a bioelectronic ion pump integrated with organoids via a custom 3D-printed holder, enabling precise ion delivery. The holder, designed with Fusion 360 and printed using biomed clear resin via Stereolithography printing technique, was positioned within a six-well plate. It offered stable support for the ion pump and allowed for easy media exchange without organoid disruption. After printing, meticulous cleaning, UV curing at 60°C, and thorough inspection ensured the holder’s quality and smooth surface finish. Sterilization eliminated potential bacterial contamination, and an O-ring secured the holder’s fit in the well plate.

The experimental protocol compromised several distinct stages. Initially, we recorded the baseline neural activity of the organoids as a reference for subsequent stages. Subsequently, we used the ion pump to deliver GABA and K^+^ ions separately to the organoids. After each delivery stage, we performed a wash step, flushing out the delivered ions and allowing the organoid to return to their baseline condition. Throughout the entire experiment, we continuously monitored the neural activity of the organoids via calcium imaging, capturing the effects of ion delivery and washout on the organoids’ neural responses.

## Acknowledgments

This work was supported by Schmidt Futures (SF857) to M.T. and S.R.S.; National Human Genome Research Institute (1RM1HG011543) to M.T. and S.R.S.; National Science Foundation (NSF2134955) to M.T. and S.R.S.. K.V. was supported by ARCS Foundation and grant T32HG012344 from the National Human Genome Research Institute (NHGRI), part of the National Institutes of Health (NIH), USA. M.R. acknowledges support from the Defense Advanced Research Projects Agency (DARPA) through Cooperative Agreement No. D20AC00003 awarded by the U.S. Department of the Interior (DOI), Interior Business Center. The content of the information does not necessarily reflect the position or the policy of the Government, and no official endorsement should be inferred and the Army Research Office (ARO) through Contract No. W911NF2210058 issued by US ARMY ACC-APG-RTP W911NF. Microfabrication was performed using equipment sponsored by the W. M. Keck Center for Nanoscale Optofluidics, the California Institute for Quantitative Biosciences (QB3), and the Army Research Office, Award No. W911NF-17-1-0460.

## Author contributions

Y.P. carried out the majority of the experiments, data analysis, and manuscript preparation. S.H. performed calcium imaging and contributed to the biological aspects of the study. C.H. was responsible for the fabrication of the bioelectronic ion pump device. H.E.S. conducted organoid imaging and calcium imaging. K.V., H.L., H.D., N.H., E.M.M., and F.N.S. provided valuable feedback and suggestions on the experimental design and manuscript. R.U., S.R.S. contributed by securing funding for the project. M.T., M.A.M.-R., and M.R. provided supervision, guidance, and conceptual ideas for the project, as well as contributed to manuscript writing, editing, and funding.

