## Supplemental Information for "Modulation of neuronal activity in cortical organoids with bioelectronic delivery of ions and neurotransmitters"

Biorxiv

Figure S1. Experimental setup

Figure S2. Calibration data for $K^+$ and GABA concentrations

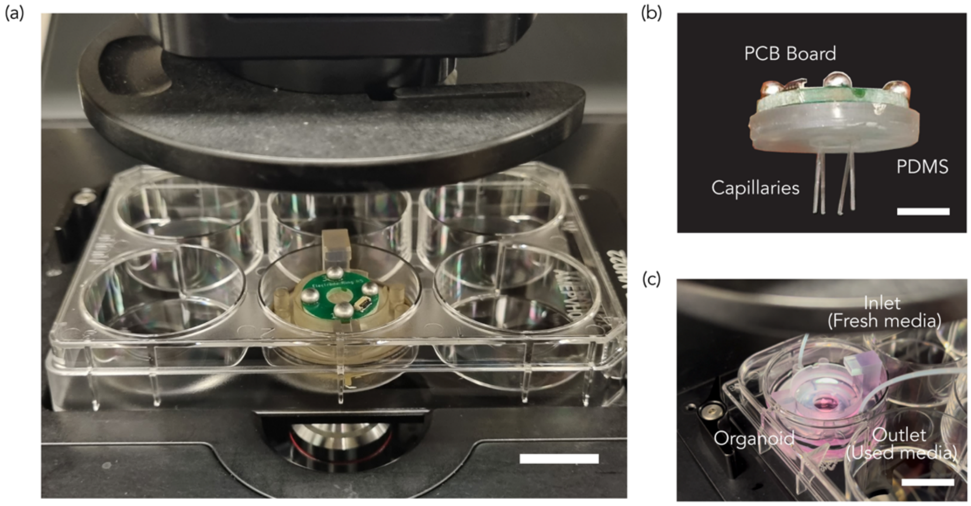


**Figure S1.** Experimental setup. (a) Experimental setup featuring a 6-well plate, with the bioelectronic ion pump device and ion pump fixed in place using a 3D-printed holder. (b) Close-up view of the bioelectronic ion pump (c) Organoid with fluidic channels for temporary ion delivery. (Scale bar: 10 mm)

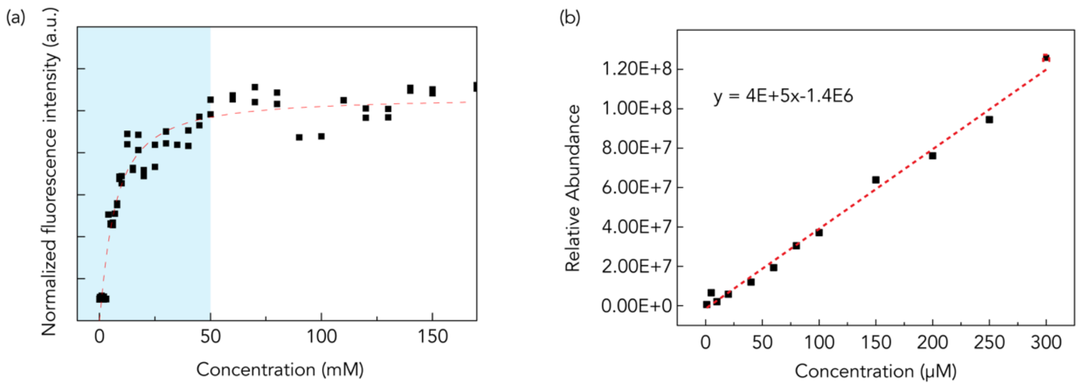


**Figure S2**. Calibration data for $K^+$ and GABA concentrations. (a) Calibration data showing the relationship between $K^+$ concentration and fluorescence intensity. (b) Calibration data displaying the relationship between GABA concentration and liquid chromatography (LC) measurements of relative abundance.
